# Plasmid copy numbers reflect distinct evolutionary strategies

**DOI:** 10.64898/2026.06.10.731345

**Authors:** Liam P. Shaw

## Abstract

Different plasmids exist at different copy numbers per cell, and there is an approximate inverse relationship between copy number and plasmid size. Two recent studies claim to quantify this relationship into a scaling law, but both the form and the interpretation of this law are contested. Here, I explore the issues with fitting a single law across plasmid diversity and suggest a consistent synthesis. First, I explore some potential problems with using sequencing-based estimates of copy number. Then, I discuss plasmid copy number through a series of case studies. I argue in favour of interpreting plasmid copy numbers not through a scaling law, but through the lens of two dominant evolutionary strategies. I argue that small plasmids which lack active segregation mechanisms will have a resulting tradeoff between plasmid inheritance and fitness cost to the host, which is responsible for their observed inverse relationship between copy number and size. In contrast, larger plasmids with active segregation mechanisms show a much weaker relationship, in line with evidence that their fitness costs are dominated by the expression of specific genes rather than their size. Where plasmids in the 20-100kb range have higher copy numbers, I argue these probably arise more from selection at the level of the host cell for plasmid-associated phenotypes (e.g. antibiotic resistance) rather than from plasmid-level selection for inheritance. These arguments are not novel, but they give good reasons for thinking that the relationship between copy number and size need not be governed by a universal constraint.

## Introduction

Plasmids contain a huge and inconvenient amount of diversity. It could have been even worse: when Joshua Lederberg first introduced the word in 1952, he intended it to be a general one for *any* extrachromosomal hereditary determinant [1]. Lederberg’s terminology was largely ignored until the 1960s, when the transfer factors capable of moving antibiotic resistance genes between different bacterial cells started to be referred to as plasmids [2]. However, other words were also used, until at a symposium in 1968 it was proposed that plasmid should be adopted as an ‘all-embracing’ term for extrachromosomal molecules that replicated autonomously [3]. As Luria wrote in the introduction to that symposium’s proceedings, ‘Bronowski once wrote that the purpose of science is the search for hidden unity in variety…[in the case of plasmids] we have lots of variety but the unity is well hidden.’

It was not obvious whether the variety could be unified. In 1972, Clowes compiled a table of available data for eleven *E. coli* plasmids and demonstrated that their copy number was proportional to their size [4]. He argued that plasmids formed two clear groups: those that occurred at ∼1-2 copies per cell and those with significantly higher copy numbers. He termed these ‘stringent’ and ‘relaxed’, after what he believed were different strengths of control mechanisms governing their copy number. Replotting his data, we can clearly see the two groups (Fig. 1). (Note the apparently anomalous position of R6K – I will return to that later.)

**Figure 1.**
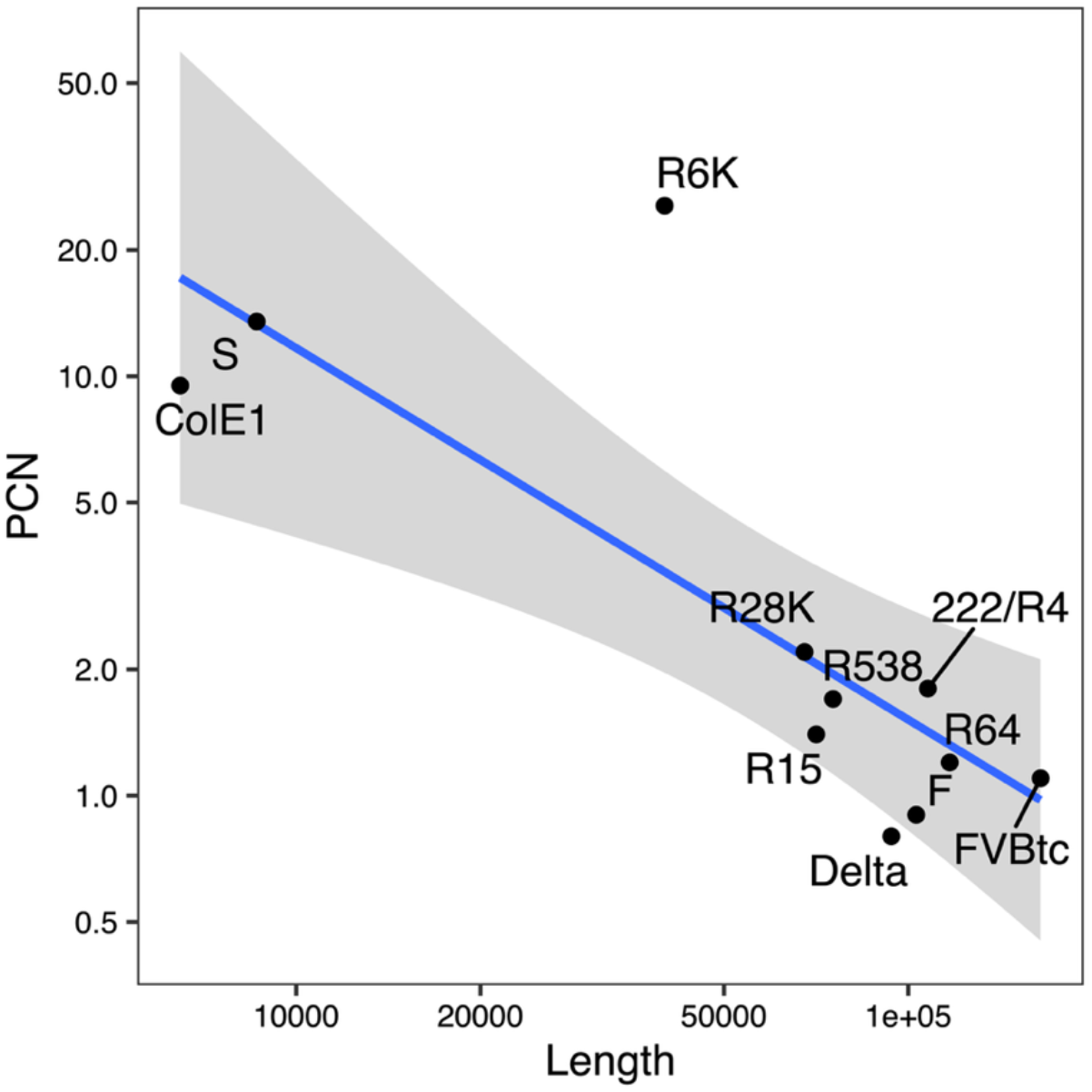
A universal scaling law between plasmid copy number and size? Data taken from Table 8 of [4]. For S, ColE1, R6K and F I have taken the mean of the two values given. Molecular weights taken from the references provided in [4] and converted to basepairs using the conversion 1bp ∼ 0.65kDa. Blue line is a linear fit log_10_ *N* = *c*+ *K* · log_10_*L*. Note that the molecular weights are used by Clowes to infer copy number from DNA ratios, so the axes are not independent measurements.

Recently, two studies have used sequencing data from thousands of plasmids to explore the relationship between copy number and size [5, 6]. Maddamsetti et al. and Ramiro-Martínez et al. both quantify the relationship into a scaling law with the general form

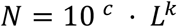

Which can rewritten for a log-log plot:

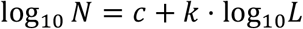

Although Clowes didn’t fit this relationship to his data, doing so produces k = −0.89, suggesting a similar approximate relationship of *N*∼1/*L* (Table 1). My point here is not to quibble over the precise value of *k*. As has been repeatedly observed prior to these papers [7–9], there is some sort of relationship between plasmid copy number and size. But what is it? And what does it mean?

**Table 1.** Values of k for a putative scaling law between plasmid copy number and size. ^#^ Value taken from the first segment of the segmented regression model for unnormalized length (Supplementary Table 3).

|  | Clowes (1972) | Ramiro-Martinez<br>(2025) | Maddamsetti<br>(2025)<sup>#</sup> |
| --- | --- | --- | --- |
| Number of<br>plasmids<br>analysed | 11 | 6,327 | 12,006 |
| $k$ | -0.89 | -0.65 | -0.96 |

In what follows, I consider how we should analyse and interpret plasmid copy numbers. My aim is to complement the unifying approach adopted by these previous papers with case studies that bring in some of the variety of plasmid biology, hoping to arrive at a synthesis.

## Methods

Maddamsetti et al. and Ramiro-Martínez et al. both calculate copy number by comparing plasmid coverage to chromosomal coverage within short-read sequencing datasets, but use slightly different methodologies. Ramiro-Martínez et al. use direct read mapping, whereas Maddamsetti et al. develop ‘Pseudoalignment and Probabilistic Iterative Read Assignment’ (pseuPIRA). Both methods give very similar results: for the 4,341 plasmids included in both datasets, the correlation between copy number estimates is extremely high (Pearson’s correlation 0.992, p<0.001). Both sets of authors also find a high correlation for their copy number estimates with data I previously published with colleagues looking at 2,292 plasmids within *Enterobacterales* [7], which is also as expected because those estimates were taken from Unicycler assembly estimates so were also based on read-mapping. Given the high concordance of these estimates, here I mainly reanalyse the larger Maddamsetti dataset, which contains 11,365 plasmids that are also present in PLSDB (94.7% of the original dataset). The Maddamsetti dataset includes n=35 plasmids from archaea (0.3% of dataset), but I do not analyse these here.

Code to reproduce results: https://github.com/liampshaw/plasmid-copy-number.

## Results and discussion

### Are sequencing-based estimates of copy number accurate?

Sequencing-based estimates of copy number are assumed to be an accurate proxy for average copy number per cell in a bacterial population. In this section, I critically consider this assumption. There are two potentially concerning factors.

First, these estimates lead to plasmids with inferred copy numbers <1. These are discarded by Ramiro-Martínez et al. (n=1000, 11.5% of initial dataset) but retained in the Maddamsetti dataset (n=2,602, 21.7% of initial dataset). Maddamseti et al. suggest various explanations [5], in particular highlighting the possibility that they can arise when the population has on average >1 chromosome per cell, as is known to happen in stationary phase [10]. Inferred copy numbers <1 are not unique to sequencing-based estimates. Clowes includes two in his dataset (Fig. 1), where values were estimated by measuring the ratio between covalently closed circular DNA and chromosomal DNA and using known molecular weights to infer copy number [4]. Values <1 could reflect a scenario of partial carriage, where some cells in the population lack the plasmid, but the plasmid is stably maintained in a (changing) sub-population through horizontal transfer. However, whatever the underlying reasons, these copy number estimates therefore do not straightforwardly reflect the average copy number in plasmid-carrying cells, complicating any inference we make about the cellular constraints that might shape copy number. Like many things in microbiology, we should remember that *in vitro* estimates may not reflect natural bacterial populations.

Second, the variance in copy number is strongly correlated with size (Fig. 2). The general trend of this relationship is expected, but it is notable that copy numbers for plasmids <10kb span at least two orders of magnitude for the same length. It is unclear to what extent this is an artefact or a genuine biological phenomenon.

**Figure 2.**
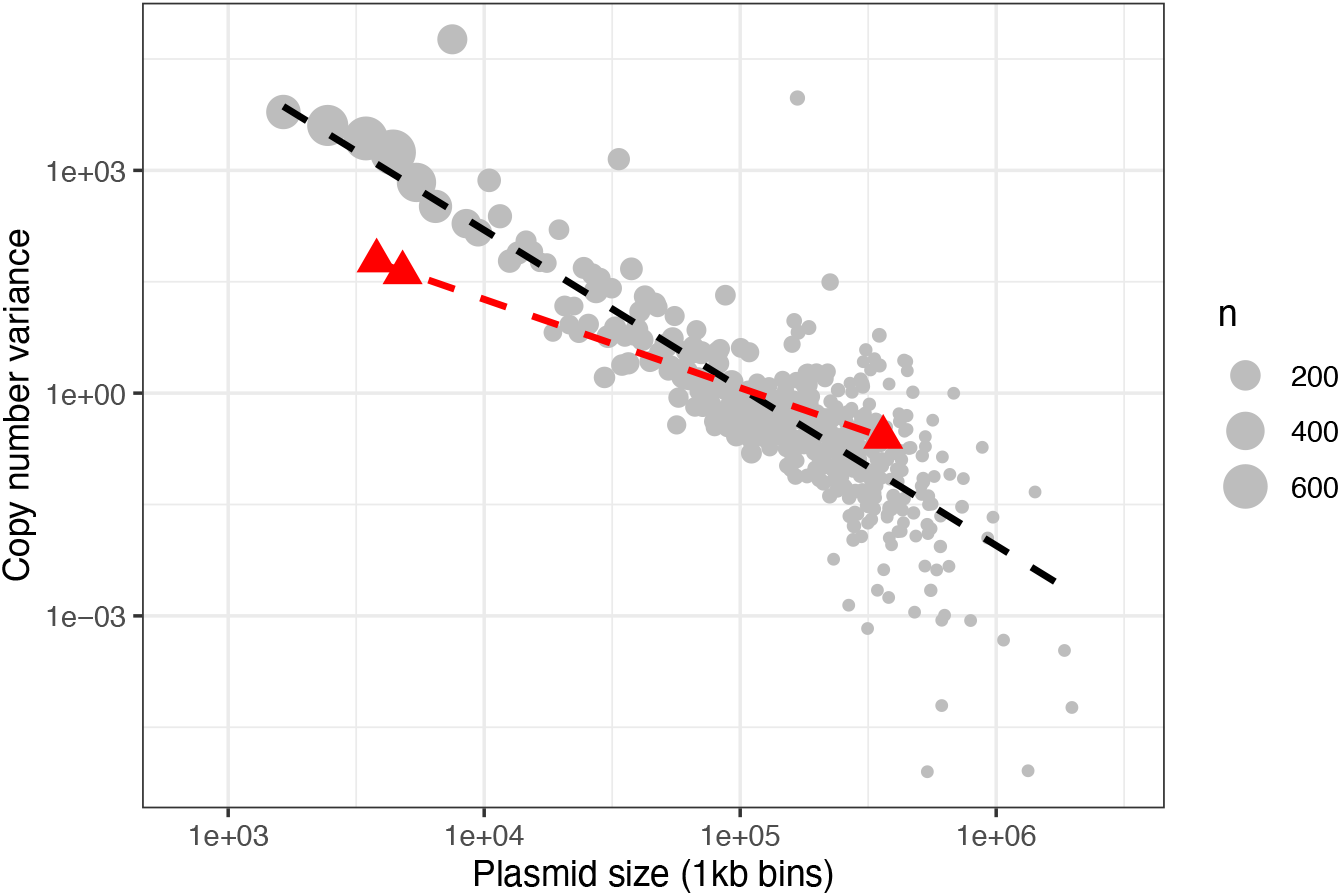
Variance in plasmid copy number is inversely correlated with plasmid size. Grey circles show variance in Maddamsetti et al. estimates [5], with plasmids grouped into 1kb bins and a linear fit shown (black dashed line). Red triangles show technical variance in sequencing-based estimates for identical plasmids in the same isolate using different DNA kits from Becker et al. [11]. Slope coefficients are -2.12 (95% CI -2.27 to -1.97) vs. -1.19 (-1.22 to -1.16) respectively.

It is known that library preparation can strongly affect sequencing-based copy number estimates. In an elegant study, Becker et al. (2016) used six DNA kits on the same isolate of a clinical strain of *Klebsiella pneumoniae* known to have three plasmids of sizes 3.8kb, 4.8kb and 382kb, then performed whole-genome sequencing [11]. The different kits did not substantively change the absolute chromosomal coverage but produced a range of copy numbers estimates for the small plasmids, with a similar inverse correlation between variance and plasmid size (Fig. 2). The 3.8kb plasmid could have an inferred PCN spanning two orders of magnitude: anything between 3 and 170 copies (median 36.0). In contrast, estimates for the 362kb plasmid had a much smaller range (1.1-1.7 copies, median 1.4). Sequencing estimates for the two smallest plasmids were much higher than those obtained with qPCR, which gave vales of 2-3x or 3-9x more abundant than the chromosome respectively; Becker et al. suggest that these qPCR results were undercounts because of supercoiling, which emphasises that other methods also have potential biases. The impact of methods for DNA purification and isolation on estimates is also supported by Plotka et al. (2017), who compared bead-beating to silica-membrane-based columns and found this led to estimates for a 4.4kb plasmid of 7.3 vs 20.5 copies [12].

However, it is also true that mutations can drive dramatic changes in copy number [13]. For example, combinations of mutations in pSC101 (9.3kb) can bring about a shift from ∼5 to ∼500 copies per cell [14]. The same plasmid may also have different copy numbers in different hosts: for example, p15a (∼5.6kb) has ∼3 copies per cell in *Vibrio natriegenes* but ∼13 in *Escherichia coli* [15]. This can even be true within a species. As an arbitrary example, consider a ColRNA plasmid found in multiple *E. coli* from different cattle farms in a single study (Table 2). This plasmid was seen in 13 isolates across three *E. coli* phylogroups (A, B1, E) with a near-identical sequence (4,671 bp, 7 variable positions). Sequencing-based estimates for its copy number ranged from 24.9-108.7 (median 34.3). But when we consider only exactly identical plasmids found in the same host strains isolated from the same farm, the copy number estimates are, reassuringly, much more similar (Table 2), supporting the claim of consistency – at least, for the case of the same host and environment with the same library preparation and sequencing methodology. It is also worth noting that stochastic variation in copy number for cells within a population may be an important route for evolutionary adaptation [16].

**Table 2.**
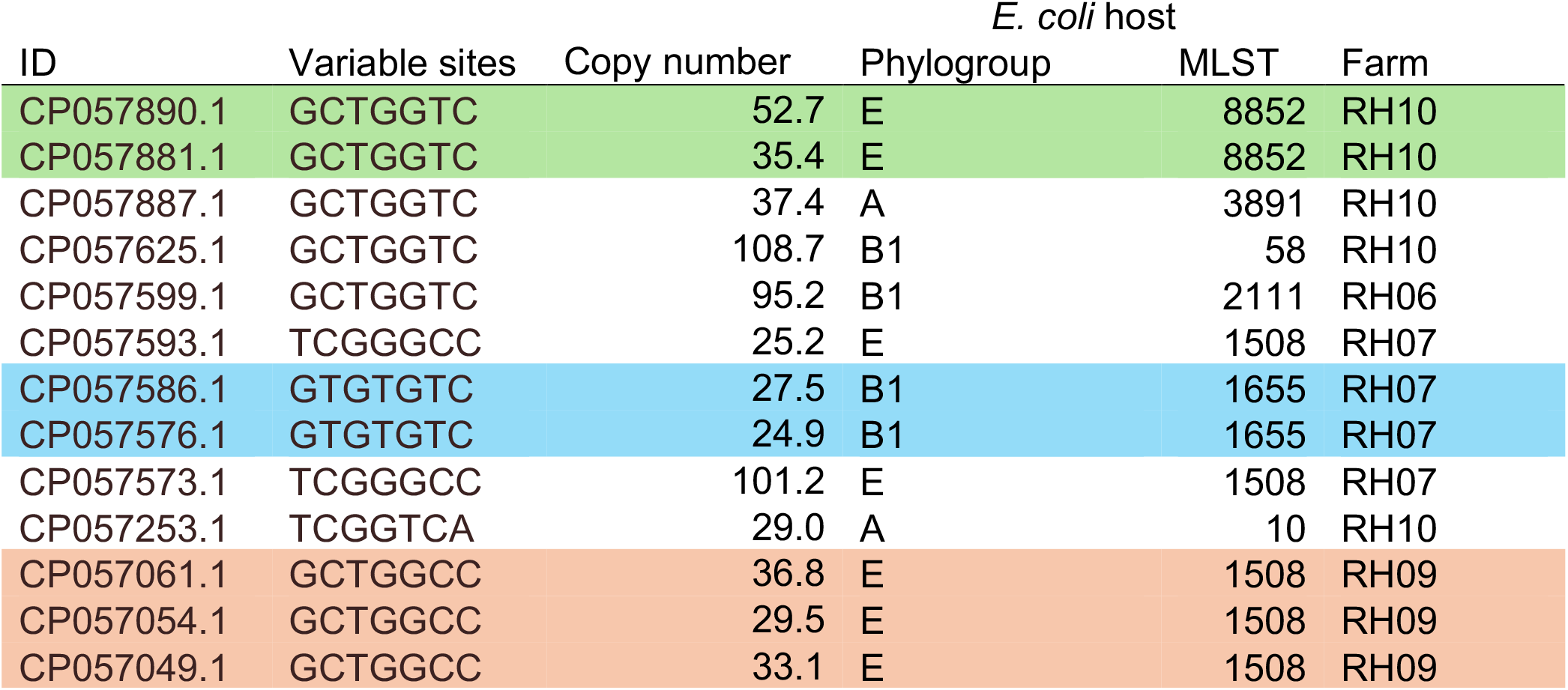
Near-identical ColRNA plasmids from livestock-associated *E. coli* have variable sequencing-based estimates for copy number. All plasmids are 4,671bp and sourced from [7]. Variable sites are shown to give a unique fingerprint (all other sites identical). MLST = multi-locus sequence type. Highlighted colours show groups of exactly identical plasmids found in the same host strains isolated from the same farm.

| ID | Variable sites | Copy number | *E. coli* host |  |  |
| --- | --- | --- | --- | --- | --- |
|  |  |  | Phylogroup | MLST | Farm |
| CP057890.1 | GCTGGTC | 52.7 | E | 8852 | RH10 |
| CP057881.1 | GCTGGTC | 35.4 | E | 8852 | RH10 |
| CP057887.1 | GCTGGTC | 37.4 | A | 3891 | RH10 |
| CP057625.1 | GCTGGTC | 108.7 | B1 | 58 | RH10 |
| CP057599.1 | GCTGGTC | 95.2 | B1 | 2111 | RH06 |
| CP057593.1 | TCGGGCC | 25.2 | E | 1508 | RH07 |
| CP057586.1 | GTGTGTC | 27.5 | B1 | 1655 | RH07 |
| CP057576.1 | GTGTGTC | 24.9 | B1 | 1655 | RH07 |
| CP057573.1 | TCGGGCC | 101.2 | E | 1508 | RH07 |
| CP057253.1 | TCGGTCA | 29.0 | A | 10 | RH10 |
| CP057061.1 | GCTGGCC | 36.8 | E | 1508 | RH09 |
| CP057054.1 | GCTGGCC | 29.5 | E | 1508 | RH09 |
| CP057049.1 | GCTGGCC | 33.1 | E | 1508 | RH09 |

To summarise, variance in copy number for small plasmids can arise from both genuine biological signal and methodological differences between studies. It seems probable that the signal we observe is some combination of these two. Although it might be hoped that averaging over large datasets should average out technical variation, without a statistical analysis controlling for experimental factors (e.g. using study as a random variable) this remains an assumption. As both sets of authors note, general patterns can still be valuable even if we should avoid over-interpreting the specific estimates for any particular plasmid. This is not a new issue: Clowes writes that we can observe important features from the copy numbers in his dataset, estimated by various methods, ‘in spite of the fact that many of these methods are subject to certain apparently uncontrollable experimental variations’ [4].

Reassuringly, evidence does suggest consistency between sequencing-based estimates and PCR-based methods. Rouches et al. found a strong correlation between copy numbers estimated from a sequencing-based method and from digital droplet PCR [17]. Furthermore, in a subsequent paper Ramiro-Martínez et al. have reported a strong correlation between copy number estimates from sequencing and qPCR at copy numbers of ∼1x, ∼10x and ∼70x (Fig. S19 of [18]; Pearson’s *r*=0.83; variation increased for higher copy numbers). Although qPCR is not a ground truth, these calibrations should give us confidence in sequencing-based estimates across the spectrum of plasmid copy number. In the rest of this analysis I will therefore assume that they represent an accurate proxy, despite their limitations.

### Interpreting the form of the scaling law

Both Maddamsetti et al. and Ramiro-Martínez et al. fitted their models to log-log data, but made different choices about their analysis methods. When a dataset has strong clustering, fitting a single linear regression can produce a highly significant overall relationship that does not hold within either of the two clusters (this is a form of Simpson’s paradox). Given that plasmids form two clusters of size and copy number, exploring the possibility of two relationships rather than one is sensible. Whereas Ramiro-Martínez et al. fitted only a single linear regression (ordinary least squares), Maddamsetti et al. used a segmented regression i.e. two linear regressions with a breakpoint, arguing that this provided evidence for a ‘biphasic’ scaling law that becomes much weaker for larger plasmids.

Reanalysing the data, a segmented regression does provide a much better fit to both datasets as judged by AIC (Fig. 3). This result is robust, holding whether we fit to the Maddamsetti dataset treating all plasmids independently (Fig. 3a) or at the level of plasmid taxonomic units (PTUs) for the Ramiro-Martínez dataset, where plasmids with copy numbers <1 were removed (Fig. 3b). This suggests that the concern in the previous section about copy numbers <1 does not lead to any qualitative difference in conclusions. Plasmid diversity has two distinct regimes, and fitting a single regression fails to capture this key feature.

**Figure 3.**
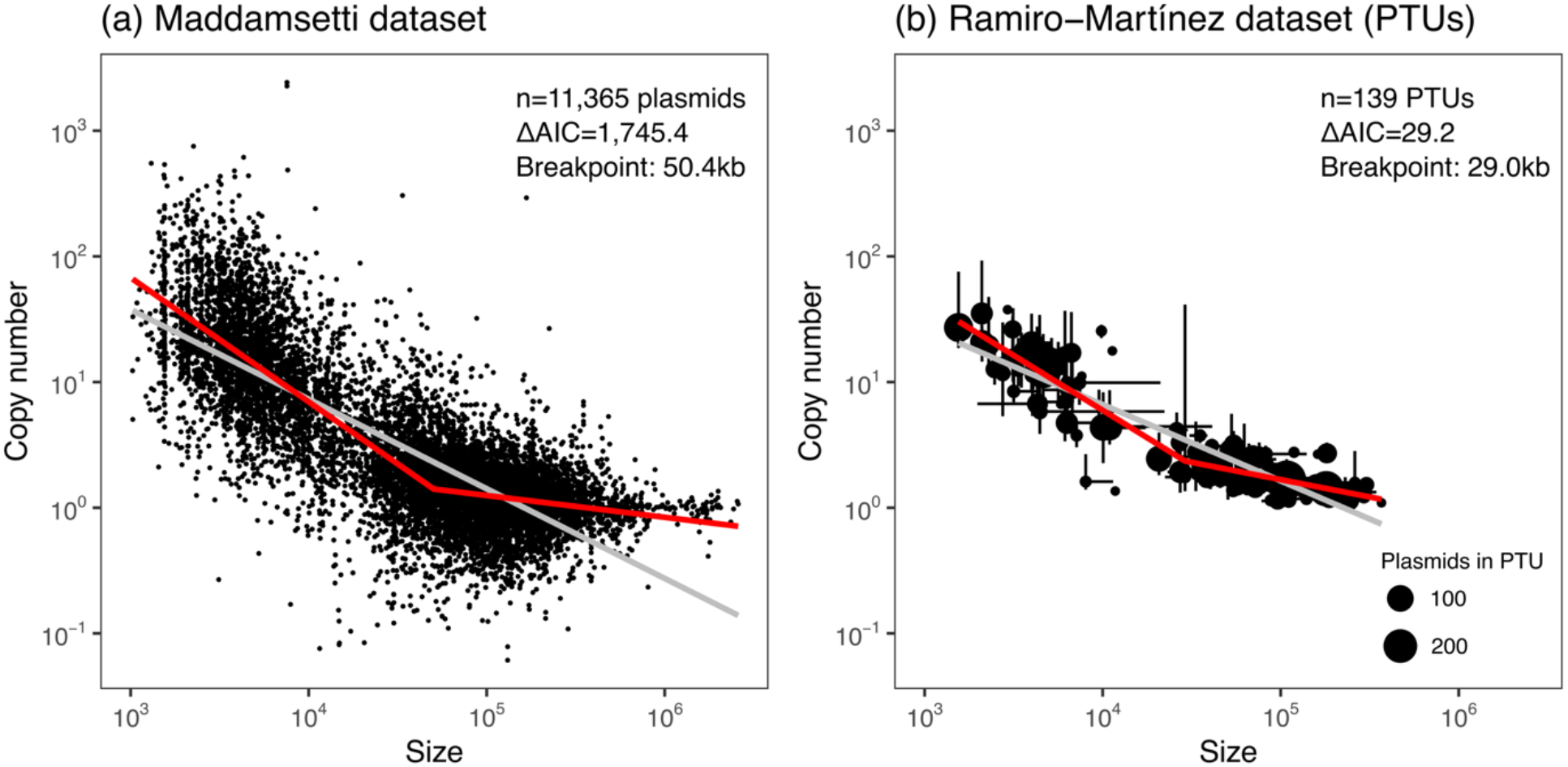
A segmented regression is a better model for plasmid copy nmber than a single scaling law. (a) Reanalysis of Maddamsetti dataset with a single (grey) or segmented (red) regression. (b) Reanalysis of Ramiro-Martínez dataset, fitting single (grey) or segmented (red) regression to 139 plasmid taxonomic units (PTUs) encompassing 4,045 plasmids (plasmids without PTU assignments excluded). Points show median values for each PTU, error bars the IQR, and size the number of plasmids within the PTU.

The breakpoint between plasmid regimes is estimated a short distance into the cluster of larger plasmids at between 29-56kb, depending on method and dataset. Interestingly, this makes the breakpoint larger than the 22.7kb that separates the Maddamsetti plasmids on size with K-means clustering (which is consistent with a ∼20kb rule of thumb for separating the bimodal distribution of plasmid lengths [19], although note that there is often confusion between the length distribution and the log-length distribution [20]). Maddamsetti et al. suggest that this breakpoint corresponds to replication strategy, marking an approximate threshold between plasmids whose replication is cell-cycle-independent vs. cell-cycle-dependent. In the next section, I discuss this interpretation.

### Interpreting the breakpoint

Clowes (1972) interpreted the two clusters of plasmids as representing two types of replication control [4]. Plasmids under ‘stringent’ control linked their replication to the cell cycle and kept their number of copies low, whereas plasmids with ‘relaxed’ control replicated more freely. The earliest mention I am aware of the terminology of ‘relaxed’ control of copy number is Rownd et al. (1966), who write ‘the control of the replication of the R-factor appears to be lax, since it was found that there are about twelve copies of the R-factor for each bacterial chromosome’ [21]. In this paradigm, the high copy number of smaller plasmids was less tightly controlled – or perhaps unregulated altogether.

This old paradigm is incorrect. All plasmids that have been studied in detail have ways of actively controlling their copy number [22]. Although the precise mechanisms are beyond the scope of this review (see [23] for a discussion), all involve some form of negative feedback, where the rate of plasmid replication is inversely proportional to copy number [9]. From a theoretical perspective, positive feedback mechanisms would be destabilizing and therefore implausible [24]. No examples are known.

Plasmid copy number is not a number that is literally encoded into the plasmid, but something that emerges from an interaction between a plasmid’s regulatory feedback control of its replication and the host environment, as discussed above for the different copy numbers of ColRNA plasmids in different *E. coli* hosts. Ramiro-Martínez et al. showed that different PTUs tend to have a characteristic copy number despite variation [6]. The key point is that the plasmid is always actively involved in this regulation: copy number is not externally determined by an external or passive mechanism, where plasmids replicate until the host makes them stop. As Summers noted in his classic textbook on plasmids in the mid-1990s, ‘despite their seductive simplicity, it is unlikely that passive mechanisms are of widespread importance in replication control’ [25]. Aside from the lack of evidence, Summers argues that host-encoded rate-limiting factors are implausible because they would need to be specific to each plasmid incompatibility group – the biochemical similarity of host functions required for replication by all plasmids argues against this.

As an example of a cell-cycle-independent plasmid, Maddamsetti et al. give the 39kb plasmid R6K (NCBI accession LT827129.1), which is smaller than their breakpoint between these lifestyles. R6K is a strange plasmid; it is anomalous in being relatively large while having high copy number [26], and is larger than the conventional ∼20kb separation between plasmid size clusters. It is also not present in the Maddamsetti dataset; R6K carries an IncX2 replicon [27] and there are no IncX2 plasmids in the Maddamsetti dataset. (The closest relative to R6K is an IncX1 plasmid (NZ_CP097723.1; mash distance <0.1 to R6K), which has an estimated copy number of 1.33 i.e. is firmly a low-copy plasmid.) More natural examples of cell-cycle-independent plasmids would have been e.g. ColE1 plasmids (typically ∼6kb), which are abundant in the Maddamsetti dataset but, unlike R6K, are clearly small plasmids.

### Size does not cleanly separate plasmids by lifestyle

The conceptual distinction between cell-cycle-independence and cell-cycle-dependence is an improvement on the incorrect distinction between relaxed and stringent control, but the heart of the problem is that size is an imperfect proxy for these two broad lifestyles. Of course, we can be confident that a plasmid lacks a particular function when the length of the genes required is larger than the entire plasmid. For example, the size of the conjugation machinery varies between plasmids, but it is always larger than 10kb e.g. for the IncW plasmid R388 it is at least 14.9kb [28], for the IncP plasmid RP4 about 15kb separated into two regions [29], and for the F plasmid around 30kb [30]. So it is safe to conclude that no plasmid <10kb is conjugative.

But when it comes to considering plasmids *above* a certain size, they can and do lack functions that other plasmids of that size tend to have, as the case of R6K shows. Large plasmids can also lose genes and their associated functions, such as conjugation [31]. Trying to fit a universal law with a single threshold runs into difficulty, because there is no critical length that cleanly divides plasmids between these two lifestyles.

This can be true even for related plasmids. For example, there are n=91 plasmids in the Maddamsetti dataset where the only detected replicon is IncFIA. These plasmids are all within the same incompatibility group, with a median copy number of 1.7 (range 0.1-10.3) and a median size of 75.3kb (13.7-274.0kb). How can we decide which of these plasmids are cell-cycle-dependent or cell-cycle-independent? For the case of the related F plasmid (NC_002483.1), its plasmid-encoded partitioning (*par*) system consists of *sopA, sopB* and *sopC*. (The F plasmid carries replicons IncFIA, IncFIB and IncFIC: IncFIB is defective [32].) Searching for *par* systems in these IncFIA plasmids reveals that plasmids lacking a *sopB* hit (searching with hmmer v3.4, hmmer.org) include those that are smallest (Fig. 4). But there are exceptions, such as a 35.5kb IncFIA plasmid that has a hit to *sopB* (NZ_CP030339.1) and some plasmids >100kb that lack a hit to *sopB* but probably have *par* systems that this crude search didn’t detect. This suggests the situation is more complicated than a single breakpoint. In fact, even this does not capture the potential complexity of copy number in these related plasmids.

**Figure 4.**
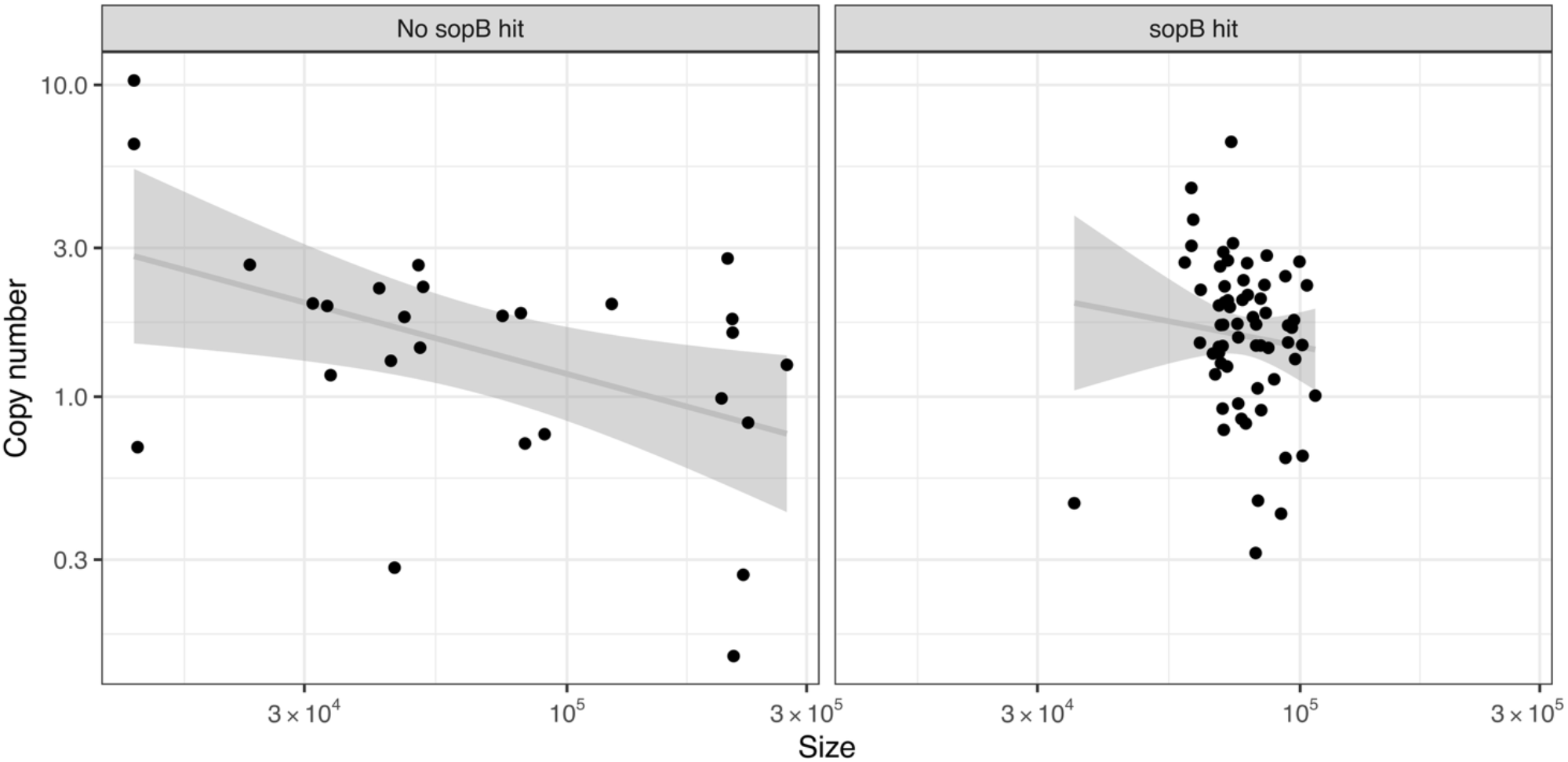
Can size alone determine cell-cycle-dependence? Plasmids with IncFIA replicon (n=91) range from 13.7-274.0kb. Searching for a hit to *sopB* (indicative of the presence of a *par* system to ensure stable inheritance) confirms that smaller plasmids tend to lack these systems, but there does not seem to be a size threshold. Grey lines show linear regression with 95% CI.

### A third lifestyle? Considering the case of ‘oligocopy’ plasmids

In the current literature, it is usually taken as a given that plasmids fall obviously into two clusters: low-copy plasmids (which are large) and high-copy plasmids (which are small). This is not strictly accurate. Alternatively, Novick (1987) divided plasmids into *three* groups based on copy number: unit-copy plasmids (1x), oligocopy plasmids (2-6x) and multicopy plasmids (>6x) [9]. Although the precise numerical division between oligo- and multicopy plasmids was arbitrary, Novick argued the conceptual division was important because some plasmids had regulatory properties that were ‘intermediate between those of unit and multicopy plasmids’. As an example he gave plasmid R1, a 97.6kb *Salmonella* plasmid with an IncFIA replicon (NZ_KY749247.1) [33]. Although R1 is usually low-copy but with copy number >1, deleting a portion of the *copB* gene which encodes CopB (a negative repressor for the *repA* promoter) results in an eightfold increase in copy number, compared to only a threefold increase in a similar 94.3kb IncFIA plasmid [34], suggesting that the regulation of copy number could be different even within the same incompatibility group.

Using Novick’s arbitrary thresholds (Fig. 5), the interquartile size ranges in the Maddamsetti dataset give some indication of the typical sizes of plasmids in these groups:

- Unit-copy plasmids (<2x): 63.8-181.9kb
- Oligocopy plasmids (2-6x): 16.8-74.6kb
- Multicopy plasmids (>6x): 3.1-6.8kb

**Figure 5.**
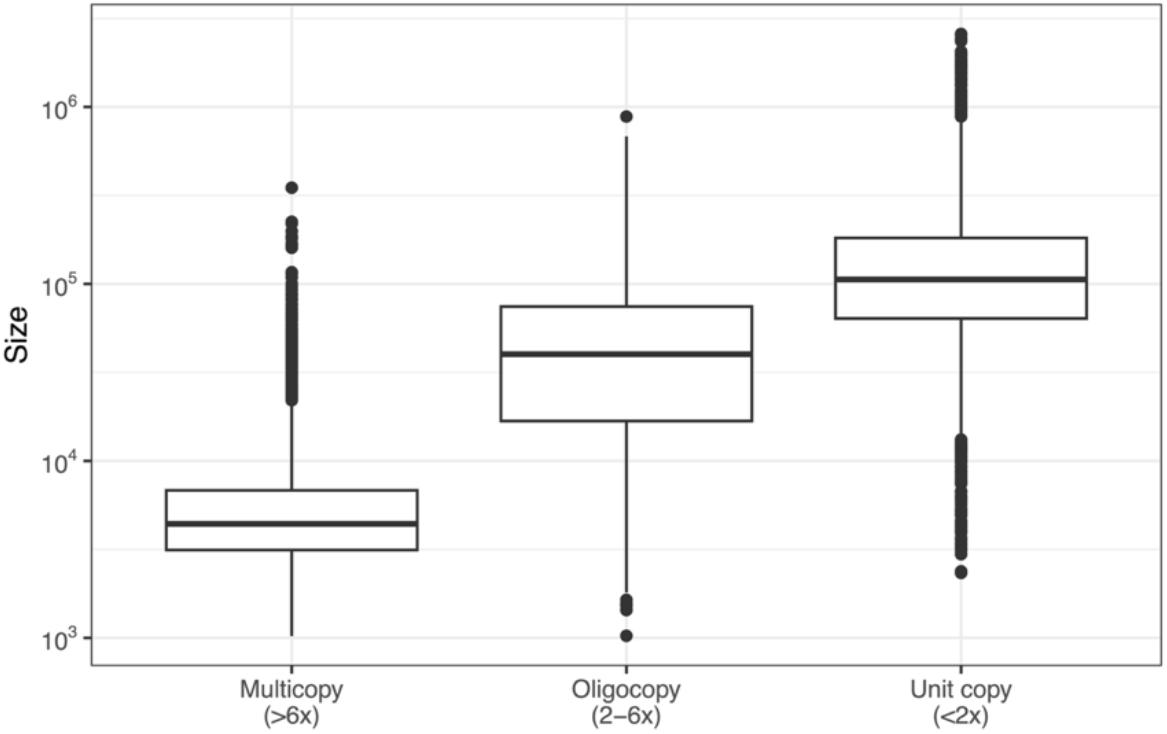
Novick’s three categories for plasmid copy number in the Maddamsetti dataset. There are 5,911 plasmids <2x, 2,279 2-6x, 3,175 >6x (52.0%, 20.1%, 27.9% respectively).

The implication of the oligocopy category is that some plasmids around the putative 30-60kb breakpoint between lifestyles (from fitting a two-regime law) may obey quite different copy number dynamics to other plasmids of the same size.

To take one example, consider IncN. In the Maddamsetti dataset there are 113 plasmids with an IncN replicon, which have a median size of 51.5kb (range 34.1-104.2kb) and median copy number 2.6. Copy numbers range from 0.4 to 18.0, suggesting a high degree of regulatory flexibility. Subsetting by the carriage of antibiotic resistance genes shows that there is an inverse relationship between copy number and size only for plasmids carrying resistance genes (Fig. 6). This suggests host-level selection for resistance to antibiotics like beta-lactams and aminoglycosides, which is encoded by dose-dependent genes carried on plasmids. In other words, we should interpret these higher copy numbers as arising not from plasmid-level selection for inheritance, but from host-level selection on plasmid-associated phenotypes.

**Figure 6.**
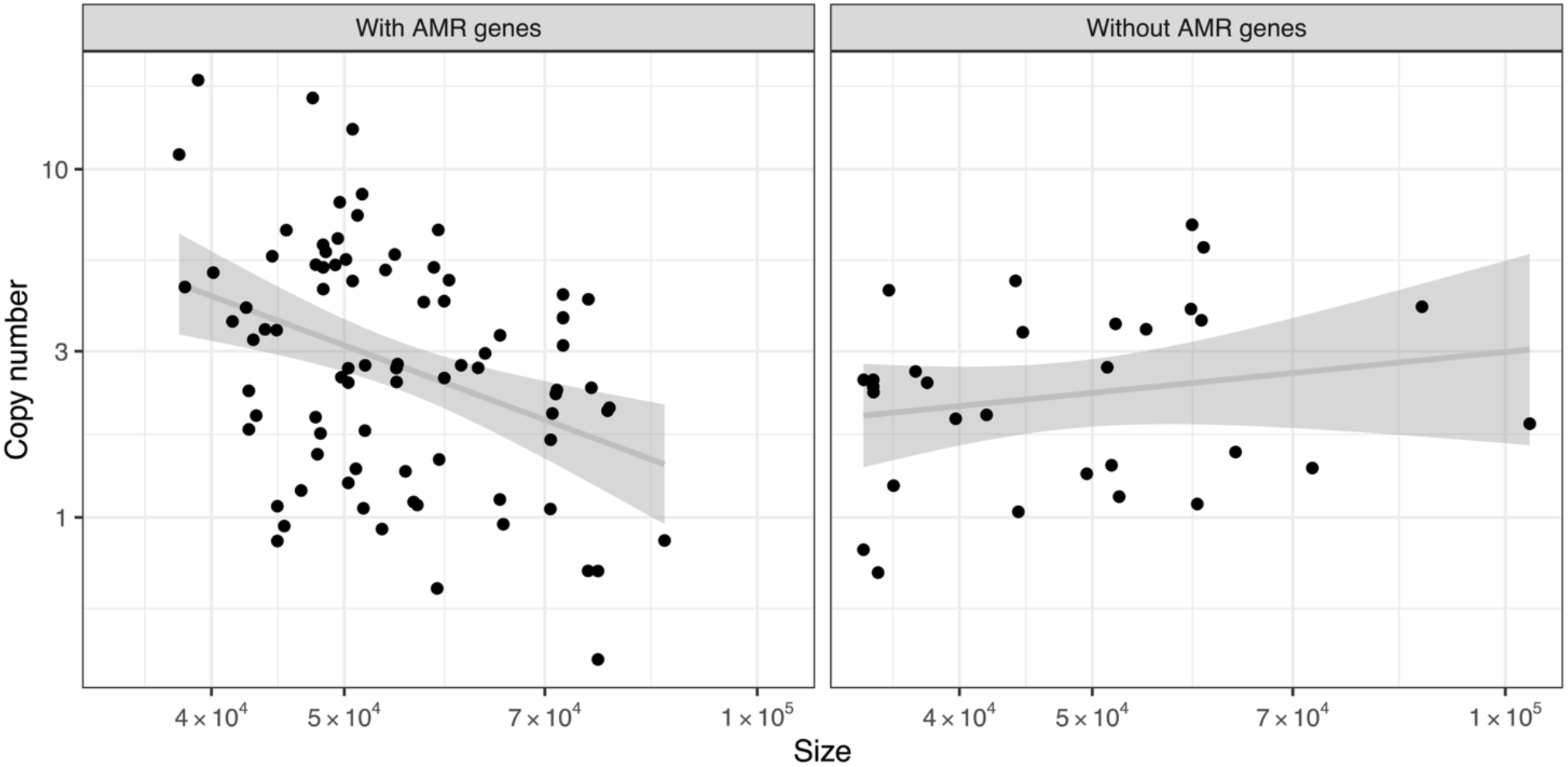
Only IncN plasmids carrying antibiotic resistance genes show a relationship between copy number and size. 113 plasmids with IncN replicons from Maddamsetti dataset, with antibiotic resistance genes detected using NCBI AMRfinderplus v4.2.7 [35].

It seems probable that oligocopy plasmids with this kind of regulatory flexibility cannot be much larger than 100kb. If we consider plasmids <100kb in 1kb size bins, those with AMR genes typically have higher copy numbers than plasmids without (65 out of 99 comparisons, mean copy number difference +1.4), supporting this interpretation of host-level selection. Although copy numbers of oligocopy plasmids are usually intermediate (in Novick’s 2-6x range), they can sometimes have much higher copy numbers due to selection at the level of the host. They therefore make a strong contribution to the relationship between copy number and size within larger plasmids. This may explain why the breakpoint from a two-regime law is larger than traditional thresholds for distinguishing small and large plasmids, and why it does not map exactly onto cell-cycle-dependence and -independence. In principle, any selection on plasmid-associated phenotypes with dose-dependence e.g. virulence or heavy metal resistance could give rise to the same pattern.

## Conclusion

Sequencing-based datasets offer a way to explore the landscape of plasmid copy number, permitting analysis of broad patterns [5, 6] as well as the case studies I have discussed here. For small high-copy plasmids less than ∼20kb (I do not think the precise value is particularly important) there is a strong but noisy inverse relationship between copy number and size, as demonstrated by both Maddamsetti et al. [5] and Ramiro-Martínez et al. [6]. I agree with Maddamsetti et al. that this relationship becomes much weaker for larger plasmids, so that a ‘biphasic’ regression with a breakpoint is much more appropriate than a single universal law. However, I have argued against over-interpreting the precise breakpoint between regimes, because this does not account for the importance of different regulatory flexibilities for different plasmids. Although we can fit approximate laws to large datasets, I believe we should not forget the underlying complexity of plasmid copy number – the diversity as well as the unity.

Claims about a universal law and the precise value of *k* bring to mind historic debates about metabolic scaling in animal biology. In that case, the ‘3/4-power law’ has now largely been abandoned in favour of an appreciation of the significant diversity involved in metabolic scaling [36]. Here, I have similarly argued for an interpretation of the available data that emphasises the diversity of plasmids; this need not be mutually exclusive with other interpretations.

I think it is important that autonomously replicating extrachromosomal molecules tend to fall into two broad classes. We call both of these ‘plasmids’, but it often makes more sense to treat them separately in terms of their distinct lifestyles or strategies, even if there is not a single length threshold that cleanly separates them. To use an analogy: most apples are smaller than oranges, but not all. However, the inevitable fuzziness of the empirical boundary *between* these fruits does not imply that we need a universal law that applies *across* them. Besides, the particular threshold from fitting a two-regime law might not be the most meaningful way to divide apples from oranges.

We can consider the two plasmid lifestyles in turn. Although small plasmids can provide advantages to their host (e.g. [37]), they seem to tend, in the limit, towards being pure DNA parasites. Although copy number can have complicated effects on host metabolism [23], at the most basic level the bioenergetic cost of any gene can be divided into three components: replication (DNA), transcription (RNA) and translation (protein). For an average bacterial gene these follow a hierarchy, with replication < transcription < translation and a one to two order of magnitude increase between replication and translation [38]. Plasmid size could increase metabolic cost to its host in a ∼1/L relationship from any of these components, but *a priori* would be expected to be dominated by translation. In support of this, small plasmids have a lower coding density as noted by Maddamsetti et al. [5], consistent with a selective pressure to reduce cost as far as possible. For high-copy molecules with low coding density, it seems possible that DNA replication would then make up a more sizeable fraction of their metabolic cost, although experiments would help to quantify this. Experiments with a ColE1 plasmid (pUC19, 2.7kb) suggest a linear metabolic burden with increasing copy number can account for the resulting reduction in cellular growth rate [17]. Another re-analysis of the Maddamsetti dataset has argued that the optimal value of plasmid copy number emerges from multi-level selection, minimising the cost function for host-level and plasmid-level fitness [39]. The general point about multi-level selection is similar to the argument by Paulsson [40]. Higher copy number therefore helps a plasmid without partitioning mechanisms to ensure its inheritance, while at the same time imposing more of a cost on the host. I interpret the approximate ∼1/L relationship that we observe for smaller plasmids (whatever the precise value of *k*) as evidence of a general tradeoff between copy number and size: the higher a plasmid’s copy number, the smaller it has to be.

The alternative lifestyle is the strategy of larger plasmids, which, in general, carry mechanisms for active segregation. The evidence is that have a much weaker relationship between copy number and size, suggesting they largely escape this tradeoff. In support of this, meta-analysis of the fitness costs of larger plasmids suggests that they are unrelated to size [41], and antibiotic resistance plasmids seem to have grown in size through the twentieth century [42]. Individual genes can have far higher translation costs than others: a recent detailed CRISPRi screen of pOXA-48 (63.6kb) across different strains showed that almost all of its cost comes from the expression of a single gene, *bla*_OXA-48_ [43]. This helps explain why the relationship with copy number is weaker in these larger plasmids: as soon as an extrachromosomal molecule does something substantial beyond encoding its own replication, its metabolic cost becomes more decoupled from its size. At the upper limit, plasmids tend towards chromosome-like properties [5]. However, not all plasmids are equal, and some plasmids <100kb can evolve reasonably high copy numbers due to regulatory flexibility (Novick’s ‘oligocopy’ plasmids, which we tend to overlook with the usual binary division between ‘low-copy’ and ‘high-copy’). I argued that these increased copy numbers likely arise from host-level selection on plasmid-associated phenotypes, rather than because of plasmid-level selection for inheritance. This is a similar distinction to the one made by Xue et al. between ‘phenotypic’ and ‘non-phenotypic’ selection [44]. We know that plasmids with antibiotic resistance genes are among the fastest-evolving and have the broadest host ranges [45]. Perhaps the capacity for regulatory flexibility may also be associated with resistance.

As a biological category, ‘plasmid’ has always contained a genuine tension between unity and variety. To me, the most arresting fact about plasmid diversity is that billions of years of evolution have been effective at keeping small plasmids small and large plasmids large, supporting the idea that there are two successful but contrasting evolutionary strategies. I do not think the available data support the conclusion that both strategies have a common constraint for the relationship between copy number and size. Most bacterial genera show a ‘valley of death’ in their plasmid size distribution, which risks being occluded if we analyse all plasmids together across bacterial diversity. (Of course, plasmids being plasmids, there are taxonomic exceptions: consider *Streptomyces* with its large, linear plasmids, but also *Clostridium, Agrobacterium* and, to a lesser extent, *Borreliella, Acinetobacter*, and *Enterococcus*.) Despite this division within plasmids being so well-known as to be a cliché that itself obscures plasmid diversity, the exact contributions that different mechanisms make to maintaining it are unclear. Intermediate plasmids face the problem of not fully exploiting either evolutionary strategy; this may be sufficient to explain their rarity, but it would be interesting to see if future evolution experiments with artificial plasmids could help us understand more precisely why.

## Acknowledgements

This article was prompted by discussion at the Microbiology Society meeting ‘Understanding and predicting microbial evolutionary dynamics’ (26-27 November 2025, Liverpool). Thanks to Craig MacLean, Rohan Maddamsetti, Lingchong You, Paula Ramiro-Martínez and Jerónimo Rodríguez-Beltrán for comments and discussion on a draft version.

